# Maximum Mutational Robustness in Genotype-Phenotype Maps Follows a Self-similar Blancmange-like Curve

**DOI:** 10.1101/2023.03.11.532236

**Authors:** Vaibhav Mohanty, Sam F. Greenbury, Tasmin Sarkany, Shyam Narayanan, Kamaludin Dingle, Sebastian E. Ahnert, Ard A. Louis

## Abstract

Phenotype robustness, defined as the average mutational robustness of all the genotypes that map to a given phenotype, plays a key role in facilitating neutral exploration of novel phenotypic variation by an evolving population. By applying results from coding theory, we prove that the maximum phenotype robustness occurs when genotypes are organised as bricklayer’s graphs, so called because they resemble the way in which a bricklayer would fill in a Hamming graph. The value of the maximal robustness is given by a fractal continuous everywhere but differentiable nowhere sums-of-digits function from number theory. Interestingly, genotype-phenotype (GP) maps for RNA secondary structure and the HP model for protein folding can exhibit phenotype robustness that exactly attains this upper bound. By exploiting properties of the sums-of-digits function, we prove a lower bound on the deviation of the maximum robustness of phenotypes with multiple neutral components from the bricklayer’s graph bound, and show that RNA secondary structure phenotypes obey this bound. Finally, we show how robustness changes when phenotypes are coarse-grained and derive a formula and associated bounds for the transition probabilities between such phenotypes.

## I. INTRODUCTION

A *genotype* describes a set of biological information. It can can be digitally encoded into a sequence of DNA or RNA, but also encoded in a more coarse-grained way, for example, in the weights of a gene regulatory network. A genotype will, in turn, map onto a *phenotype*, which is a biologically observed output, trait, or behaviour, in what is known as a genotype-phenotype (GP) map [1–3]. Examples include 4 letter RNA sequences and 20 letter protein sequences that can be mapped to their physical folded states, and gene-regulatory networks, which can, for example, be described by Boolean networks [4] where a set of weights represent the gene interaction strengths. Because many mutations are effectively neutral there will typically be many more genotypes than phenotypes. The set of all genotypes that map to a given phenotype is called a neutral set, or sometimes also a neutral network. It plays an important role in shaping the way that novel variation arises in evolutionary dynamics. Properties of neutral sets have been extensively studied [1–24]. A number of key features of neutral sets shared across GP maps [2, 3, 18]. For example, the size of the neutral sets for different phenotypes typically vary over many orders of magnitude, with a small fraction of the phenotypes taking up the majority of genotypes. Such phenotype bias can strongly affect evolutionary outcomes [14, 17, 24–30].

Another important shared trait is that neutral sets are typically highly connected by point mutations due to a high average mutational robustness, meaning that they are likely to be fully connected, or percolate. This property hugely enhances the probability that a neutral set can be traversed by single mutational steps, allowing a much larger set of alternate phenotypes to be accessible than one could reach from a single genotype. In this way, enhanced robustness can lead to enhanced evolvability, which is the ability to discover new phenotypes [9, 31]. In some cases, the neutral set is split into smaller component networks which are disconnected, for example due to biophysical constraints [7, 10, 32]. Then it is not the full neutral set, but rather each neutral component that percolates and is easily traversed via point mutations.

The property that we will focus on in this paper is the mutational robustness *ρ*_*p*_ of a phenotype *p*, defined as the average probability that a single character mutation of a genotype mapping to phenotype *p* does *not* change the phenotype *p*. Typically larger neutral sets have higher robustness. For the RNA sequence to secondary structure GP map, it has been shown that the distribution of robustness found upon random sampling of sequences accurately predicts the distribution of robustnesses for functional or non-coding RNAs found in nature [33], although for very short strands, naturally occurring RNA are marginally more robust [24]. In other words, for this system, the structure of GP map appears to largely determine the mutational robustness found in nature. Thus studying these more abstract mathematical features of the GP map may directly lead to predictions about naturally occurring phenotypes.

To study *ρ*_*p*_ in a mathematically convenient way, we will use the language of graphs. The entire set of *k*^*ℓ*^ possible sequences in a GP map with input sequences of length *ℓ* drawn from an alphabet of *k* characters is representable as a generalisation of a hypercube graph called a *Hamming graph H*_*ℓ,k*_. Each sequence maps onto a vertex, and two vertices are connected by an edge in *H*_*ℓ,k*_ only if the corresponding sequences differ by a single character. The neutral set of all genotypes mapping to phenotype *p* define a vertex set *V* (*G*_*p*_) on a vertex-induced subgraph (or neutral set) *G*_*p*_ in *H*_*ℓ,k*_. In other words each vertex represents one of the genotypes in the neutral set. Similarly, the edge set *E*(*G*_*p*_) is defined as the set of all edges between vertices in *G*_*p*_, and represents genotypes that are connected by neutral mutations. The robustness can now be defined as the average degree of *G*_*p*_ divided by *ℓ*(*k* − 1) such that it is normalised to 0 ≤ *ρ*_*p*_ ≤ 1. Using the relationship between the average degree and the edge-to-vertex ratio, we have a graph-theoretic definition of robustness *ρ*(*G*_*p*_) of the neutral set *G*_*p*_:

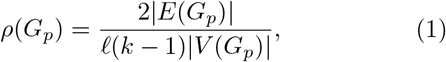

In other words, this definition of the robustness scales with the average number of edges per vertex, normalised so that it is equal to the fraction of possible mutations in the neutral set that are neutral. Relatively high phenotype robustness is important because to good approximation, if *ρ*_*p*_ *>* 1*/*(*ℓ*(*k* − 1)), then the graph *G*_*p*_, or equivalently, the neutral set or neutral component, should percolate [18], which greatly facilitates neutral exploration.

A naive first guess at the robustness predicts that the probability of a mutation being neutral should scale as

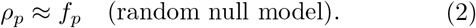

where the frequency *f*_*p*_ ≡ |*V* (*G*_*p*_) |*/k*^*ℓ*^ is equal to the probability of obtaining *p* when choosing a genotype at random. This scaling holds when genotypes are completely uncorrelated as they would be in a random null model where genotypes are randomly attributed to phenotypes under the constraint that the neutral set sizes are kept fixed. For the typically strong phenotype bias observed in GP maps, this scaling implies that many phenotypes will have a robustness that is too small to allow neutral set graphs to percolate.

However, empirical studies of robustness have consistently shown a log-linear relationship with the frequency of a phenotype:

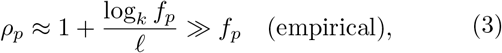

for a wide range of GP maps in biology, as well as for similar input-output maps in computer science, and physics [2, 3, 15–23]. The empirically measured robustness is orders of magnitude higher than what is predicted by the random null model. There must therefore be strong correlations between the genotypes mapping to a particular phenotype [18]. One important biological consequence of this empirically measured higher robustness is that typically, a large fraction of all neutral set graphs in a GP map should percolate, leading to enhanced evolvability.

These empirical results also raise an interesting question that will be the main focus of this paper, namely *What is the upper bound on robustness, and how close are real GP maps to this bound?* Here, we prove, by applying concepts from coding theory harking back to the 1960s [34, 35], that a particular type of graph called *bricklayer’s graphs* [36] are maximally robust.

We then derive explicit expressions for the maximum robustness of bricklayer’s graphs, which by extension gives a maximum on the possible robustness of neutral sets. These expressions are closely related to the well known sums-of-digits function from number theory. The dominant term scales as 1 + log_*k*_ *f*_*p*_*/ℓ*, which resembles the empirical scaling of eq. (3). Next, we numerically confirm that many individual neutral components for the RNA sequence to secondary structure GP maps and the hydrophobic-polar (HP) protein folding GP map introduced by Dill [37] can attain the bricklayer’s graph bound exactly.

We prove that if a full neutral set is made up of several neutral components, then the maximal attainable robustness will be below the maximum given by a bricklayer’s graph. By deriving a new property of the sums of digits function, we are able to calculate a lower bound on the robustness of a neutral set made up multiple connected neutral components. We show numerically that the RNA GP map obeys this lower bound.

Lastly, we consider the coarse-graining of phenotypes. We show how robustness and transition probabilities change when multiple phenotypes are combined into compound phenotypes, and use our equations to explain some, at first glance, non-intuitive results about the robustness of compound phenotypes.

## II. BRICKLAYER’S GRAPHS HAVE MAXIMUM ROBUSTNESS

The term “bricklayer’s graph” was coined by Reeves *et al*. [36] in an interesting paper where they studied the principal eigenvalue of the subgraph, which is a different measure of phenotype robustness because it also takes into account population structure at steady state [38]. The name arises because they are constructed by repeatedly adding an adjacent vertex in the Hamming graph in a manner resembling the process of laying bricks. While these particular graphs have a much older origin in coding theory, we will use this more recent nomenclature throughout this work. Bricklayer’s graphs are formally defined as follows:

### Definition II.1.

*A* ***bricklayer’s graph*** *G*_*n,k*_ *is an induced subgraph of a Hamming graph H*_*ℓ,k*_ *containing* |*V* (*G*_*n,k*_) | = *n vertices* {0, 1, …, *n* − 1} *such that* (*i, j*) ∈ *E*(*G*_*n,k*_) *if the base-k representations of i and j differ in exactly one digit*.

From our definition of the robustness (1), it follows that maximising robustness for a given number of vertices *V* (*G*) (the size of graph *G* that represents the neutral set) is equivalent to maximising the number of edges of the equivalent graph. Finding the subgraph *G* with a fixed number of vertices of a Hamming graph *H*_*ℓ,k*_ that maximises the number of edges (and robustness) is equivalent to minimising the “edge boundary” of the subgraph, i.e. minimising the number of edges {*u, v*} which connect a subgraph vertex *u* ∈ *V* (*G*) to a vertex outside the subgraph *v* ∈ *V* (*H*_*ℓ,k*_\*G*). This is known as the “edge-isoperimetric problem” for the Hamming graph. This problem is connected to coding theory because, for example, it was effectively proven in [34] and [35] that if the vertices of a bricklayer’s graph (which they referred to as a vertex numbering which maximizes “connectedness,” similar to robustness) represent the set of *k*-ary sequences that map to a particular codeword being transmitted, then this set of sequences minimises the number of singlesite mutations that would cause an incorrect transmission of the codeword, averaged over all the sequences mapping to that codeword. This is akin to maximising mutational robustness, which measures the average number of point mutations that do *not* change the phenotype, in biological systems. More specifically, Harper [34] showed that bricklayer’s graphs attain the maximum bound for the *k* = 2 case, and Graham [39] and Hart [40] calculated the exact value of the bound for *k* = 2, namely |*E*(*G*)| ≤ *S*_2_(*n*), where 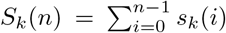 is formulated in terms of *s*_*k*_(*i*), the sum of all digits in the base-*k* representation of the integer *i*. We will call *S*_*k*_(*n*) the *sums-of-digits function*.

Importantly for this study, Lindsey [35] generalised the work of Harper [34] to prove that bricklayer’s graphs (not necessarily uniquely) attain the maximum bound for all *k* ≥ 2 although he did not calculate the value of the bound. No other graph can have a robustness higher than a bricklayer’s graph, although some graphs may attain the same robustness.

## III. ROBUSTNESS OF BRICKLAYER’S GRAPHS

### A. Exact Robustness/Number of Edges

To exactly calculate the maximum robustness, we first need to prove a theorem about the maximum number of edges of a bricklayer’s graph:

#### Theorem III.1.

*A bricklayer’s graph G*_*n,k*_(*V, E*) *with n vertices has* 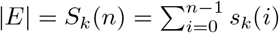 *edges, where s*_*k*_(*i*) *is the sum of all digits in the base-k representation of the integer i, and S*_*k*_(*n*) *is the sums-of-digits function*.

*Proof*. See Appendix A. □

This theorem generalises the proof by Graham [39] and Hart [40] for all *k* ≥ 2, and improves on the bound given by Squier *et al*. [43].

To calculate the upper bound on robustness, we therefore need to work out the properties of the sums of digits function. As far back as 1940 Bush [44] already showed that the asymptotic behaviour (for large n) of the sums of digits function scaled as 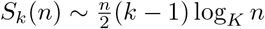. An exact analytical form for *k* = 2 was given by Trollope in 1968 [45] and later generalised by Delange in 1975 for all *k* [46] as:

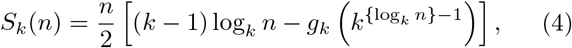

where {*x*} denotes the fractional part of *x*, and

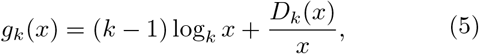

where *D*_*k*_(*x*) is the *Delange function* (using the modified definition in ref. [47]) given by

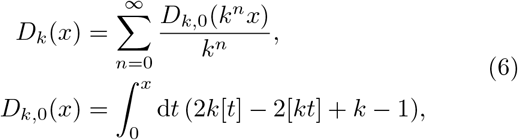

where [*x*] is the integer part of *x*. For *k* = 2, the Delange function *D*_*k*_(*x*) is the same as the continuous everywhere, differentiable nowhere Takagi function first described in 1903 [41]. The fractal Takagi function is sometimes called the Blancmange function, because it resembles the blancmange dessert. It has many applications, including to mathematical analysis, probability theory, and number theory [48] The general sums-of-digits function has also interesting connections to many fields, and in particular to number theory. For example, Delange [46] famously showed that the Fourier series coefficients {*c*_*n*_} of *g*_*k*_ (*k*^{*x*}−1^) (which is periodic in *x* with a period of one) are defined by

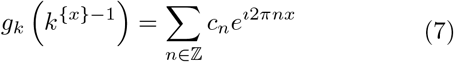

with

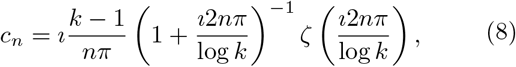

are linked to *ζ*, the Riemann zeta function.

Combining our expression for phenotype robustness (1) with theorem III.1 and eq. (4) provides an expression for the maximum phenotype robustness for a neutral set of size *n*_*p*_ = |*V* (*G*_*p*_)|:

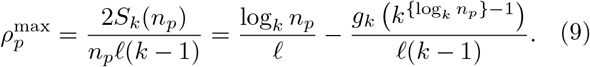

This upper bound is optimal because we can always construct a bricklayer’s graph with *n*_*p*_ vertices. The maximum robustness is plotted in Figure 1, and exhibits the expected Blancmange-like self-similar form. Since the frequency *f*_*p*_ = *n*_*p*_*/k*^*ℓ*^, the first term reproduces the empirically observed scaling of eq. (3), namely *ρ*_*p*_ *≈ ℓ*^−1^ log_*k*_ *n*_*p*_ = 1 + log_*k*_ (*f*_*p*_)*/ℓ*, which upperbounds the exact robustness in Fig. 1.

**FIG. 1.**
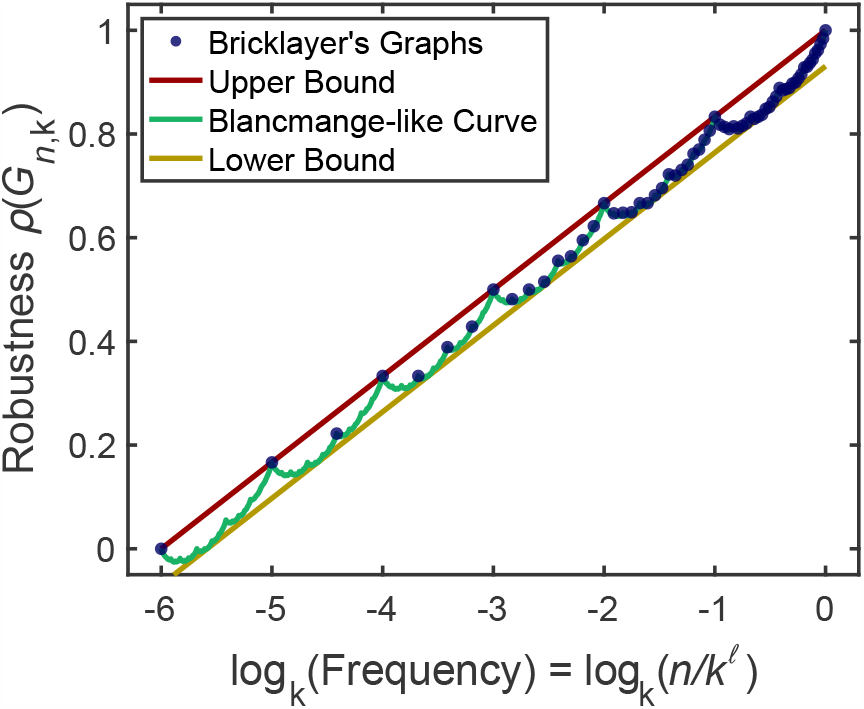
A linear-log plot of the bricklayer’s graphs’ robustness 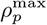 for *n* vertices, given by eq. (9), versus frequency (number of vertices *n* divided by *k*^*ℓ*^), where *ℓ* = 6 and *k* = 2. Each blue dot denotes a possible neutral set size. The green line denotes the continuous everywhere but differentiable nowhere “blancmange-like curve” (here *k* = 2, so one component of this line is exactly equivalent to the Tagaki curve [41]) that is given by the continuous *n*_*p*_ version of eq. (9), corresponding to 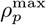. The upper and lower bounds on 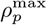, given by eq. (11), are also plotted. The upper bound is equivalent to the simple form *ρ*_*p*_ = *ℓ*^−1^ log *n*_*p*_ = 1 + log (*f*_*p*_)*/ℓ*. Plots like this, containing the exact maximum robustness as well as the upper and lower bounds, can be generated with our free, open-source web tool RoBound Calculator [42].

### B. Bounds on Neutral Set Robustness

It would be useful to find simpler expressions to bound the maximum robustness. Indeed, Galkin and Galkina [47] specify the bound

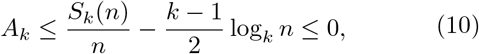

in terms of a constant *A*_*k*_ that only depends on the alphabet size *k*. This implies that the robustness of a bricklayer’s graph, or equivalently the maximum robustness of a neutral set of size *n* from eq.(9) is bounded below and above by

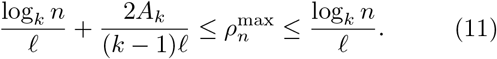

In other words, the empirical scaling (3) is a strict upper bound to the robustness, and it also provides a lower bound up to terms that scale with 𝒪 (1*/ℓ*). We have created a free, open-source web tool called RoBound Calculator using Google Colaboratory which can generate, for specified *k* and *ℓ*, generate exact values and continuous interpolations of the bricklayer’s graph robustness 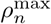, as well as the upper and lower bounds from eq. (11). RoBound Calculator is available at ref. [42], and several example plots from this tool are provided in Appendix C.

There is no short formula to calculate *A*_*k*_, but Galkin and Galkina [47] have found an exact, though fairly involved, algorithm to determine *A*_*k*_. For *k* = 2, *A*_2_ = log_4_ 3−1 ≈ −0.2075; for *k* = 3, *A*_3_ = log_3_ 2−1 ≈ −0.3691 and for *k* = 4, *A*_4_ = (3*/*4) log_2_ 5 − (9*/*4) ≈ −0.5086 (biologically relevant, as DNA/RNA have *k* = 4). Additional biologically relevant values of *k* can be calculated using their algorithm. For instance, proteins have *k* = 20 amino acids comprising their primary sequence, and we can use the algorithm to find that 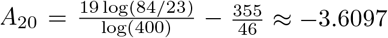. Moreover, as *k* → ∞,

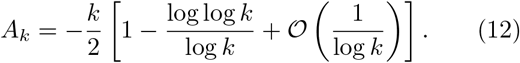

Finally, note that the correction term 2*A*_*k*_*/*((*k* − 1)*ℓ*) that appears on the left hand side of the eq. (11) is typically quite small compared to the scale of the typical values of *ρ*_*p*_ found for neutral components of these systems. For instance, this correction term has the value −0.0283 for RNA12, and −0.0226 for RNA15. eq. (11) is therefore typically a tight bound (see also Fig. 1). Thus, the maximum robustness can be quite reasonably approximated by simple 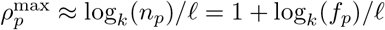 form.

## IV. NEUTRAL COMPONENTS OF BIOLOGICAL GP MAPS CAN ATTAIN THE BRICKLAYER’S GRAPH BOUND

We next turn to the question of how close the neutral components of real GP maps are to the upper bound (9). We study the RNA (length 12 and 15) secondary structure models with all 4 nucleotides (ACUG) as well as the HP protein folding models (length 24, and 5 × 5 lattice). The RNA and HP simulation data were obtained from prior work [18].

A neutral set for a phenotype may not fully percolate, but it can be broken into multiple neutral components of various sizes that do percolate [18] (though some phenotypes may have only a single component). By definition, a neutral component has no single mutation connections to any other neutral component of that same phenotype’s neutral set.

In Figure 2, the robustness values of each neutral component in the RNA12, RNA15, HP24, and HP5 × 5 models are plotted against the logarithm of the number of vertices in that neutral component. For all of the systems studied, many neutral components over several orders of magnitude of phenotype frequencies *f*_*p*_ exactly attain the same robustness as bricklayer’s graphs. Typically this is more common for smaller than for larger neutral components. In the HP model GP maps, the size of the largest neutral components that still reach the bricklayer’s graph line have fewer vertices than the same for RNA secondary structure GP maps. This is likely due to the architecture of the GP maps themselves; it has been shown recently [49] that neutral components are often modular in that they consist of highly packed clusters of vertices which are then connected to other clusters by a smaller set of linking vertices. We speculate that in the HP model GP map there may be higher modularity, leading to less-than-maximally robust neutral components above a lower threshold. It is also worth mentioning the RNA and HP systems examined here are fairly small in length because it is computationally quite expensive to exhaustively check the neutral components for much larger sequence lengths.

**FIG. 2.**
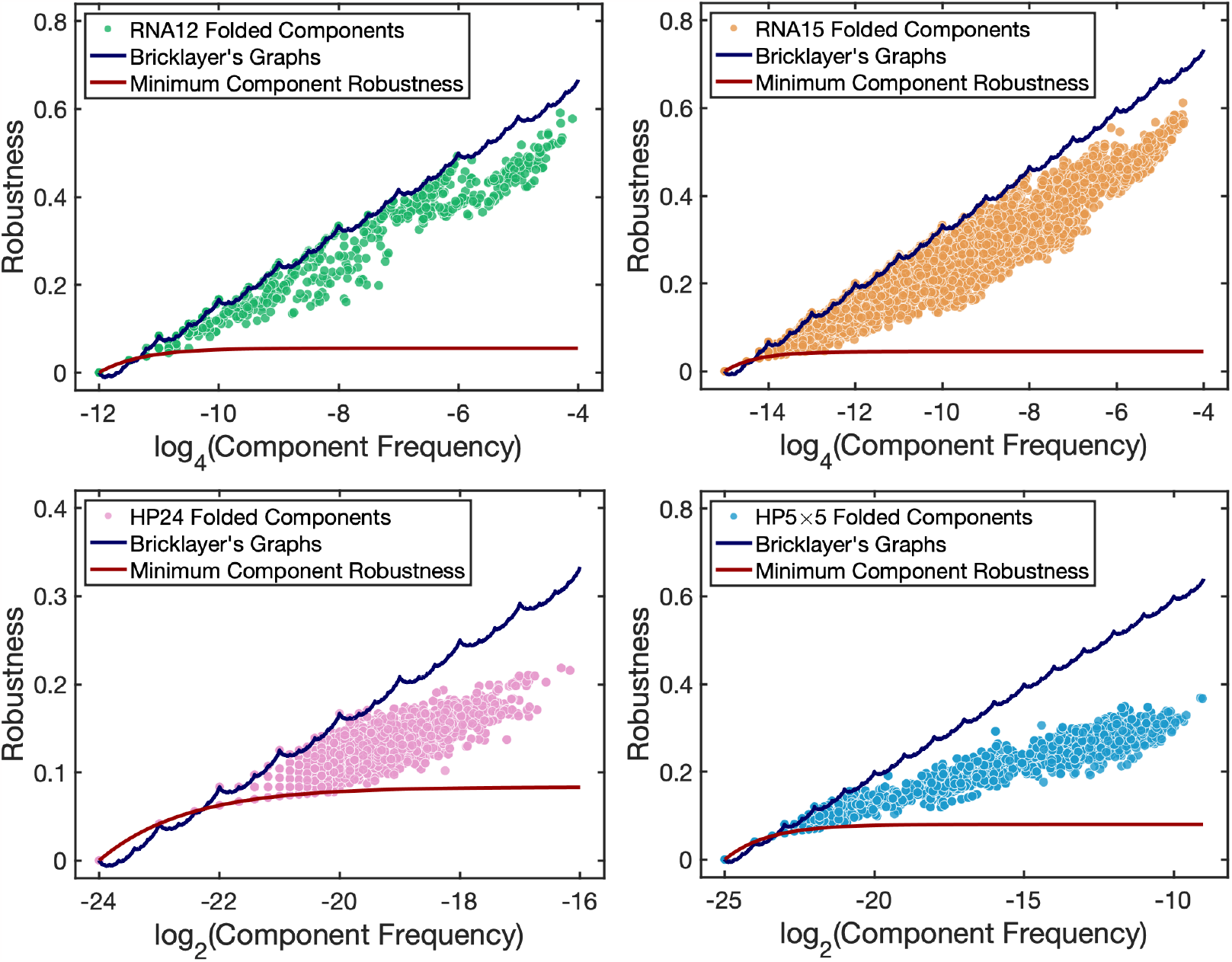
The robustness of every component of every folded phenotype for both the RNA and HP GP maps (of various lengths) is plotted against the frequency (fraction of vertices *f* = |*V* (*G*) | */k*^*ℓ*^ in the entire Hamming graph) alongside the continuous interpolation of the maximum robustness curve from eq. (9)) (blue line). The minimum robustness of a neutral component, eq. (13) is given by the (red line). The data for these curves can be generated by using the RoBound Calculator [42] tool we have introduced. The natural GP maps all contain neutral components which attain the bricklayer’s graph bound (as well as some very low robustness components that attain the minimum bound). The unfolded (trivial) phenotype is omitted from each of these plots. The minimum robustness appears to be larger than the bricklayer’s graph line for low frequencies; but, this only happens for non-integer values of the number of vertices. Of course, any graph will have an integer number of vertices; in all of those cases, the bricklayer’s graph robustness will be greater than or equal to the minimum robustness.

Figure 2 also depicts the minimum robustness of each fully connected neutral component *G*_*i*_:

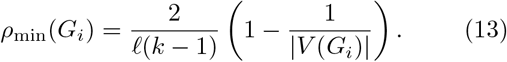

The above formula follows from the fact that a neutral component, by definition, is connected, and the minimum number of edges in a connected graph with *n* vertices is *n* − 1. This minimum value can be attained by many graphs, including the path graph *P*_*n*_ and star graph *K*_1,*n*_. This is also the robustness that individual phenotypes/outputs have for the 1D Edwards-Anderson spin glass input-output map [23]. Note that the null-model for robustness, *ρ*_*p*_ ≈ *f*_*p*_ includes many disconnected components, and so can be much lower than eq. (13), which holds for a fully connected component. The RoBound Calculator tool [42] we have introduced also calculates and plots the *ρ*_min_ curve as well.

The strict upper bound (9) is close to empirical scaling (3) observed for many GP maps, and quite far from the naive scaling *ρ*_*p*_ ≈ *f*_*p*_. Since real robustness cannot be higher than the bound, this suggest that, at least on the scale given by *f*_*p*_, real GP maps exhibit a robustness that is close to the maximum value attainable. This empirical observation raises interesting theoretical questions because the upper bound is not imposed by the specific properties of an individual GP map (biological or not); rather, the robustness is bounded above due to very general mathematical properties of the Hamming graph underlying the GP map.

## V. ROBUSTNESS OF FULL NEUTRAL SETS AND THE BRICKLAYER’S BOUND

A bricklayer’s graph is maximally dense in its edge-to-vertex ratio; it must certainly be fully connected. Therefore, if a neutral set is not fully connected, and thus broken down into neutral components, then the full phenotype robustness *ρ*_*p*_ must deviate from this optimum, even if the robustness of each of its components attains the optimal bound.

To calculate a bound on *how much* phenotypes broken down into neutral components deviate from the optimal robustness, we consider a neutral set that has *n* vertices and is split into *m* neutral components. If each neutral component is maximally robust (as many of the RNA/HP neutral components are), then each robustness would be

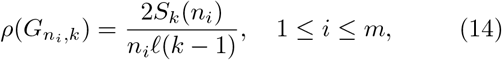

where 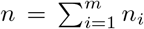, with *n*_*i*_ being the number of vertices in the *i*-th neutral component. In other words, the total robustness *ρ*_*p*_ for this specific case of a neutral set made up of bricklayer’s graph components is simply a frequency weighted sum of the robustnesses of the individual neutral components:

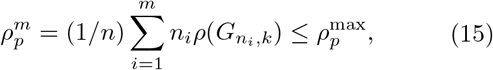

where the last inequality simply follows from the fact that 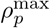 is the strict upper bound on robustness, which can only be obtained when the entire neutral set only has one single component.

To calculate a more accurate bound, we first prove an interesting property of the sums-of-digits function *S*_*k*_(*n*), generalising the proof by Graham [39], who proved the following for *k* = 2, which we now prove for general *k*:

### Theorem V.1.

*For k nonnegative integers* {*n*_1_, *n*_2_, …, *n*_*k*_} *obeying* 0 ≤ *n*_1_ ≤ *n*_2_ ≤ … ≤ *n*_*k*_, *the following property of the sums-of-digits function holds:*

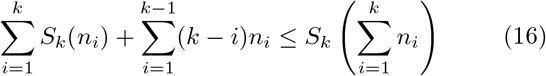

*Proof*. See Appendix A. □

Theorem V.1 is not only an interesting property of the sums-of-digits function which may be useful in coding theory, but it also be used to provide, for the specific case that there are exactly *k* neutral components, a tight bound

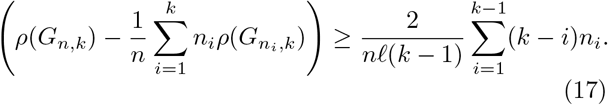

on the difference between the maximum phenotype robustness for a fully connected neutral set of size *n*, given by eq. (19) and 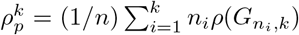, the robust-ness of of a neutral set consisting of *k* bricklayer’s graphs as neutral components. A further generalisation and discussion of this formula is provided in Section VI.

The assumption that the neutral components are bricklayer’s graphs is not necessary, however. Weakening this assumption to assume that each neutral component has an arbitrary topology simply weakens the tightness of the bound. For an arbitrary neutral component 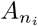 with *n*_*i*_ vertices, 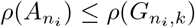, so

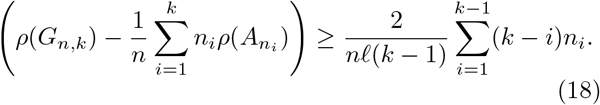

It is important to note that this inequality has been proven to hold only when the number of neutral components is less than or equal to *k*. Indeed, it does hold perfectly for the biological RNA neutral component/phenotype robustness data from our dataset from ref. [18]. In Figure 3, each plot point represents a phenotype, and the vertical axis coordinate is given by the log of the left hand side of eq. (18), which is (the log of) the difference in the optimal number of edges and the actual number of edges for that phenotype. The horizontal axis coordinate is given by the log of the right hand side of eq. (18), which is a theoretical bound computed from the frequencies of the neutral components for that phenotype. The left plot in Figure 3 shows actual RNA12 phenotype data, and the right plot shows RNA 15 phenotype data. Green plot points have *k* = 4 or fewer neutral components; it is for these phenotypes that the theoretical bound rigorously holds. The biological data support the theory if all green plot points are *above or on* the dashed 1:1 diagonal line, as this would indicate that the inequality is valid; indeed that is the case.

**FIG. 3.**
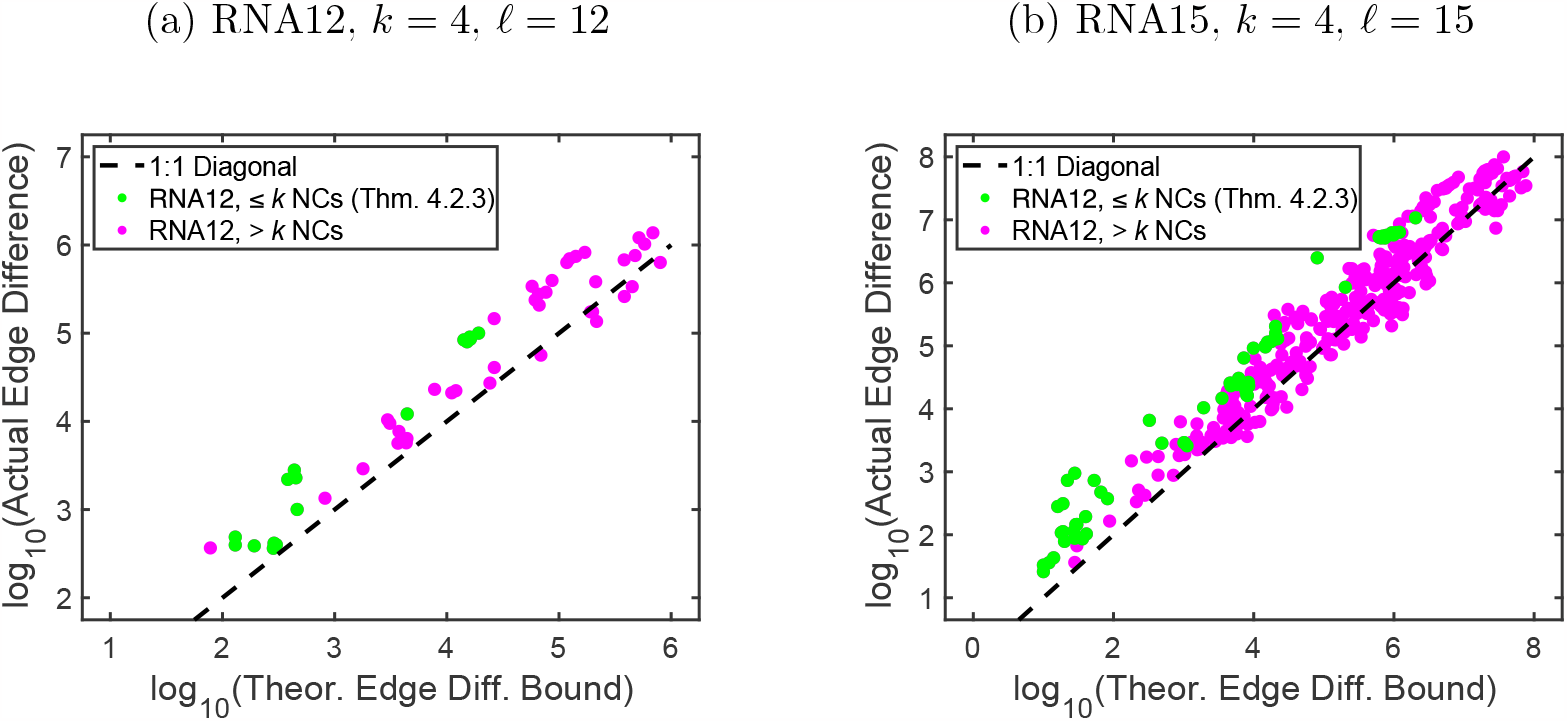
Plot of left hand side (ordinate) and right hand side (abscissa) of eq. (18) for real phenotype data for RNA12 and RNA15 from our dataset from ref. [18]. Green plot points represent phenotypes with ≤ *k* = 4 neutral components; the theoretical bound in eq. (18) rigorously holds for these phenotypes. Magenta plot points have *>* 4 neutral components; despite the fact that the bound should not rigorously hold for such phenotypes, it still does seem to approximately hold for many phenotypes, or at least many plot points lie close to the dashed line.

Most of the phenotypes in the RNA12 and RNA15 secondary structure GP maps do not have ≤ *k* = 4 neutral components, however. Also in Figure 3, we have plotted, in magenta, the values of the left and right hand side of eq. (18) for phenotypes with *> k* = 4. The theoretical bound seems to hold even for most of the cases outside the range for which it is proven, but it begins to fail for sufficiently large phenotypes.

Here, we discussed the process by which many neutral components are combined into one larger phenotype; these neutral components are not connected to each other (by definition), so the phenotype robustness is simply a frequency weighted average of the component robustnesses that will be necessarily lower than the maximum possible achievable robustness for a single-component phenotype. With this being said, in practice, an evolving population will typically be confined to a component, as double neutral mutations are typically quite rare. Therefore, even though the robustness of the phenotype is lower, the robustness experienced in shorter timescales by the population may often be closer to the bricklayer’s graph bound.

## VI. ROBUSTNESS OF COARSE-GRAINED OUTPUTS/PHENOTYPES

In discussing neutral set and neutral component topologies for natural systems, one should keep in mind that the definition of a phenotype can vary depending on which biological property one is interested in. Working out exactly how to map genotypes to phenotypes in biological systems can be difficult due to questions of how a phenotype is defined. For RNA structures, for example, one may be interested in a specific secondary structure, as we are here, or in a more restricted phenotype, such as a certain tertiary structure. Conversely, one may be interested in a broader class of secondary structures. or in a different property, such as a certain catalytic function. It is therefore interesting to ask how robustness changes if one zeroes in on different levels of description of a phenotype.

In this section, we will study the robustness of a phenotype and transition probabilities between phenotypes that have been generated from the union of multiple neutral sets. We refer to this process of merging neutral sets of different phenotypes as “coarse-graining” of phenotypes. Tackling this question requires a more generalised approach than was used in the previous section because different phenotypes typically have nonzero transition probabilities between each other, unlike neutral components which have zero transition probabilities (on single mutations).

We illustrate our approach for RNA secondary structure using a coarse-graining method from Giegerich *et al*. [50], who defined a set of new “abstract shape” RNA secondary structures which ignore fine details of the stem and loop lengths and nesting. There may, in fact, be good biological reasons for doing this coarse-graining if one thinks that function or structure is not that sensitive to small changes in the full secondary structure. Another reason for our interest in this coarse-graining scheme comes from the work of Dingle *et al*. [29], who showed that the frequency with which non-coding or functional RNA abstract structure appears in the Rfam database [51, 52] of non-coding or functional RNA could be remarkably well predicted over 5 orders of magnitude by the frequency by which these structures appear upon random sampling of sequences. This phenomenon illustrates how biases in the arrival of variation, which are mediated through the GP map, can dramatically affect evolutionary outcomes [14, 24].

In coarse-grained GP maps, the value of the robustness is determined by the level of coarse-graining as well as by the underlying neutral set topologies. To study this problem we will first derive analytic formulas which follow from the underlying graph theory. We then present numerical results on coarse-grained RNA GP maps and explain a qualitative changes in robustness with phenotype coarse-graining. We conclude by deriving a critical transition probability that would be needed for two coarse-grained phenotypes to maintain “high” robustness when coarse-grained together.

### A. Robustness and Transition Probabilities for Coarse-Grained Phenotypes

As before, we consider a GP map whose genotypes consist of sequences of length *ℓ* drawn from an alphabet of *k* characters. The genotype space is the Hamming graph *H*_*ℓ,k*_, and phenotype neutral sets are induced subgraphs of *H*_*ℓ,k*_. The *i*-th phenotype’s neutral set *G*_*i*_ (assuming 1 ≤ *i* ≤ *N*_*p*_, where *N*_*p*_ is the total number of phenotypes) is an induced subgraph of *H*_*ℓ,k*_. Once again, we let *V* (*G*) denote the vertex set of a graph *G, E*(*G*) denote the edge set of *G*, and we additionally define *E*(*G*_*i*_, *G*_*j*_) = *E*_*H*_ (*G*_*j*_, *G*_*i*_) to denote the set of edges induced in graph *H*_*ℓ,k*_ by union *V* (*G*_*i*_) ∪ *V* (*G*_*j*_) which are neither elements of *E*(*G*_*i*_) nor *E*(*G*_*j*_), where we have taken both *G*_*i*_ and *G*_*j*_ for *i* ≠ *j* to be induced subgraphs of *H*_*ℓ,k*_. Precisely, *E*(*G*_*i*_, *G*_*j*_) = {{*u, v*} ∈ *H*_*ℓ,k*_ | *u* ∈ *V* (*G*_*i*_) ∧ *v* ∈ *V* (*G*_*j*_)}.

As we have defined before, the robustness of the *i*-th phenotype is defined as

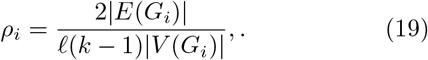

In other words it is proportional to the ratio of edges to vertices in the neutral set graph. The transition probability that a single point mutation in the genotype leads to a change from phenotype *i* to phenotype *j* is defined by

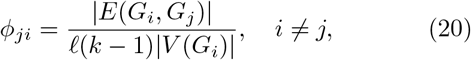

which is again the average number of links per node between the the two distinct neutral sets *i* and *j*. Note that *ϕ*_*ji*_ |*V* (*G*_*i*_) | = *ϕ*_*ij*_ |*V* (*G*_*j*_) |. If we define the diagonal terms *ϕ*_*ii*_ ≡ ρ_i_ then there needs to be an additional prefactor of 2 since the connections are between nodes of the same neutral set, and so must be counted twice.

#### 1. Robustness of Coarse-Grained Phenotypes

We first derive a general formula for robustness of coarse-grained phenotypes. Let *S* be the set of phenotype indices that indicate which phenotypes are being coarse-grained into a new neutral set *G*_*S*_. The vertex set of *G*_*S*_ is the union of all vertices

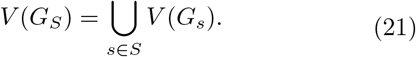

The edge set of *G*_*S*_ includes all edges in each individual neutral set as well as the edges joining the neutral sets:

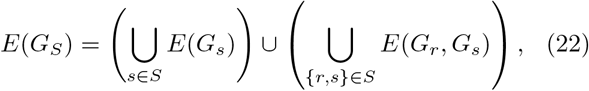

where (*r, s*) ∈ *S* denotes an ordered pair of elements of *S* such that *r ≠ s*. It follows that the robustness *ρ*_*S*_ of coarse-grained phenotype *S* is

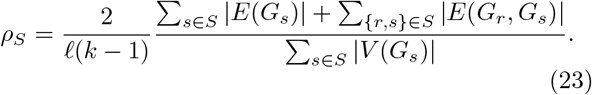

Using eq. (19) and the normalised phenotype frequency *f*_*i*_ = |*V* (*G*_*i*_) | */k*^*ℓ*^, we can rewrite the coarse-grained robustness in terms of familiar biological parameters

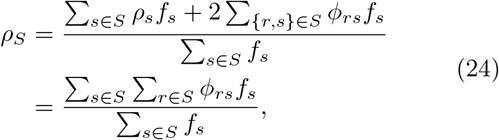

where {*r, s*} ∈ *S* denotes an unordered pair of elements of *S* such that *r* ≠*s*, and in the last step we have used *ϕ*_*ss*_ = *ρ*_*s*_. It is easy to check that, Lif *S* = {1, 2, …, *N*_*p*_} is the set of *all* phenotypes, then 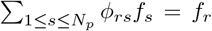 (given the definition of *ϕ*_*rs*_), so *ρ*_*S*_ = 1 as expected. The intuition behind eq. (24) is that the robustness of the coarse-grained phenotype includes terms that come from the frequency weighted sum of the robustnesses of the original phenotypes, as was the case for combining neutral components in eq. (15), plus a term that adds in the contribution from transition probabilities between the combined phenotypes. If there are many transitions between them, then the combined robustness will be higher because these are now also classed as an additional contribution to robustness.

#### 2. Transition Probabilities between Coarse-Grained Phenotypes

We now calculate a general formula for transition probabilities between coarse-grained phenotypes. Let *S* and *T* be two non-overlapping sets of phenotype indices that indicate which phenotypes are being coarse-grained into two coarse-grained neutral sets *G*_*S*_ and *G*_*T*_, respectively. The set of edges *E*(*G*_*S*_, *G*_*T*_) joining *G*_*S*_ and *G*_*T*_ is the union of all sets of edges that adjoin every pair of (non-coarse-grained) phenotypes, where within each pair one element is picked from the constituent phenotypes of *S* and the other is picked from constituent phenotypes of *T*. It follows that

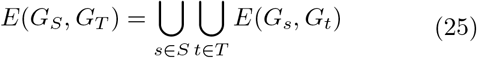

It now follows that the transition probability *ϕ*_*T S*_ from the *S*-th coarse-grained phenotype to the *T* -th coarse-grained phenotype (assuming *S* and *T* have no overlap) is

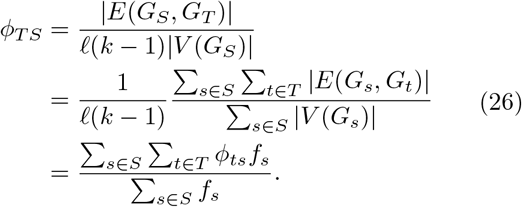

We can now see from eq. (24) and eq. (26) that the coarse-graining procedure takes on the same functional form, which is represented graphically in Figure 4.

**FIG. 4.**
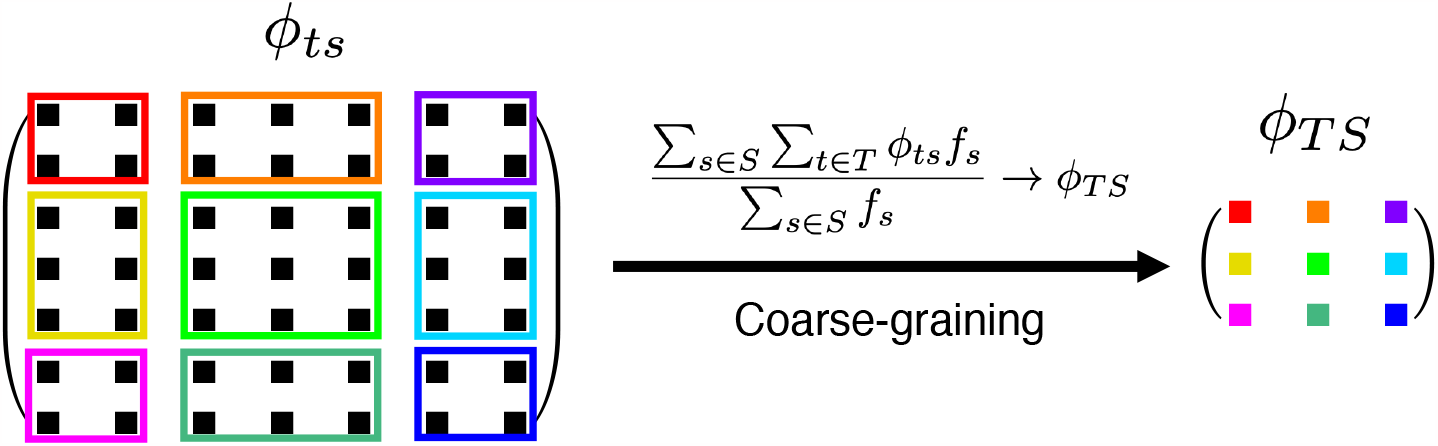
Schematic diagram of phenotype coarse-graining on the transition matrix *ϕ*_*ts*_ → *ϕ*_*TS*_, which includes transition probabilities (off-diagonals) and robustness (diagonals). If two non-overlapping sets of original phenotypes are coarse-grained into two new coarse-grained phenotypes *T* and *S*, then the transition probability from coarse-grained phenotype *S* to coarse-grained phenotype *T* is given by *ϕ*_*TS*_, calculated in eq. (26), which involves taking a frequency-weighted sum over the transition probabilities *ϕ*_*ts*_ between the original non-coarse-grained phenotypes that comprise *T* and *S*.

The process of coarse-graining neutral components into a phenotype neutral set, handled in Section V, is a specific case of the more general process of coarse-graining phenotype neutral sets, but in the former case, all transition probabilities between neutral components *ϕ*_*ji*_ = 0 since neutral components are not connected to each other, by definition. We now examine coarse-graining in in the RNA secondary structure GP map. The RNAshapes program [53] merges RNA phenotypes at different “levels” of coarse-graining by progressively ignoring more and more layers of detail in the stem and loop lengths and nesting structure of the folded RNA oligonucleotides. The “dot-bracket” structure is the actual RNA secondary structure phenotype; Level 1 of coarse-graining ignores some details of the dot-bracket structure and combines similar phenotypes into the same abstract phenotype; the Level 2 structures include further coarse-graining, and so forth. There are 5 possible levels of coarse-graining. A schematic of the RNA coarse-graining process is shown in Figure 5. In Figure 6, we present results in which RNA secondary structure GP maps have robustness values calculated for these various levels of coarse-graining, performed using the RNAshapes tool.

**FIG. 5.**
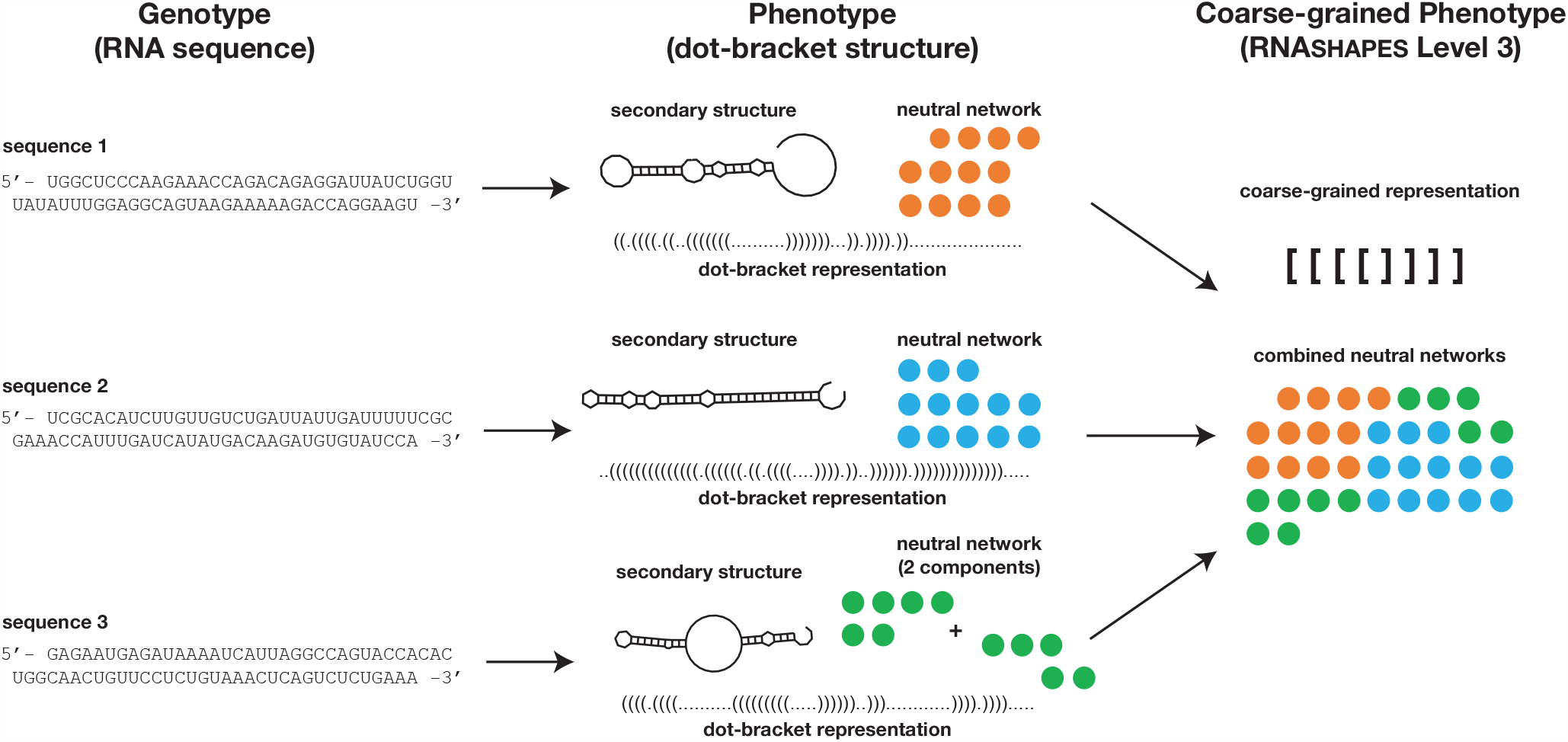
Demonstration of coarse-graining for RNA secondary structures. Genotypes (RNA sequences) map to secondary structure phenotypes represented as dot-bracket structures; the RNASHAPES program [53] yields results similar to the ViennaRNA program [54]. RNASHAPES then progressively computes coarse-graining at different “levels,” increasingly ignoring nesting of secondary structure topological features. In this figure, 3 example sequences of length *L* = 70 are are shown to map to their dot-bracket phenotypes. A cartoon of a neutral set (not actual size) has been drawn for each phenotype; network properties like robustness can be calculated for each phenotype. At at a higher level of coarse-graining (here, Level 3), multiple dot-bracket phenotypes all map onto the same coarse-grained phenotype. Accordingly, the coarse-grained phenotype has a neutral set comprised of the individual neutral sets of the original phenotypes. Note that in practice there are many more secondary structures beyond those shown that map to this same coarse-grained phenotype; the analysis above is schematic.

**FIG. 6.**
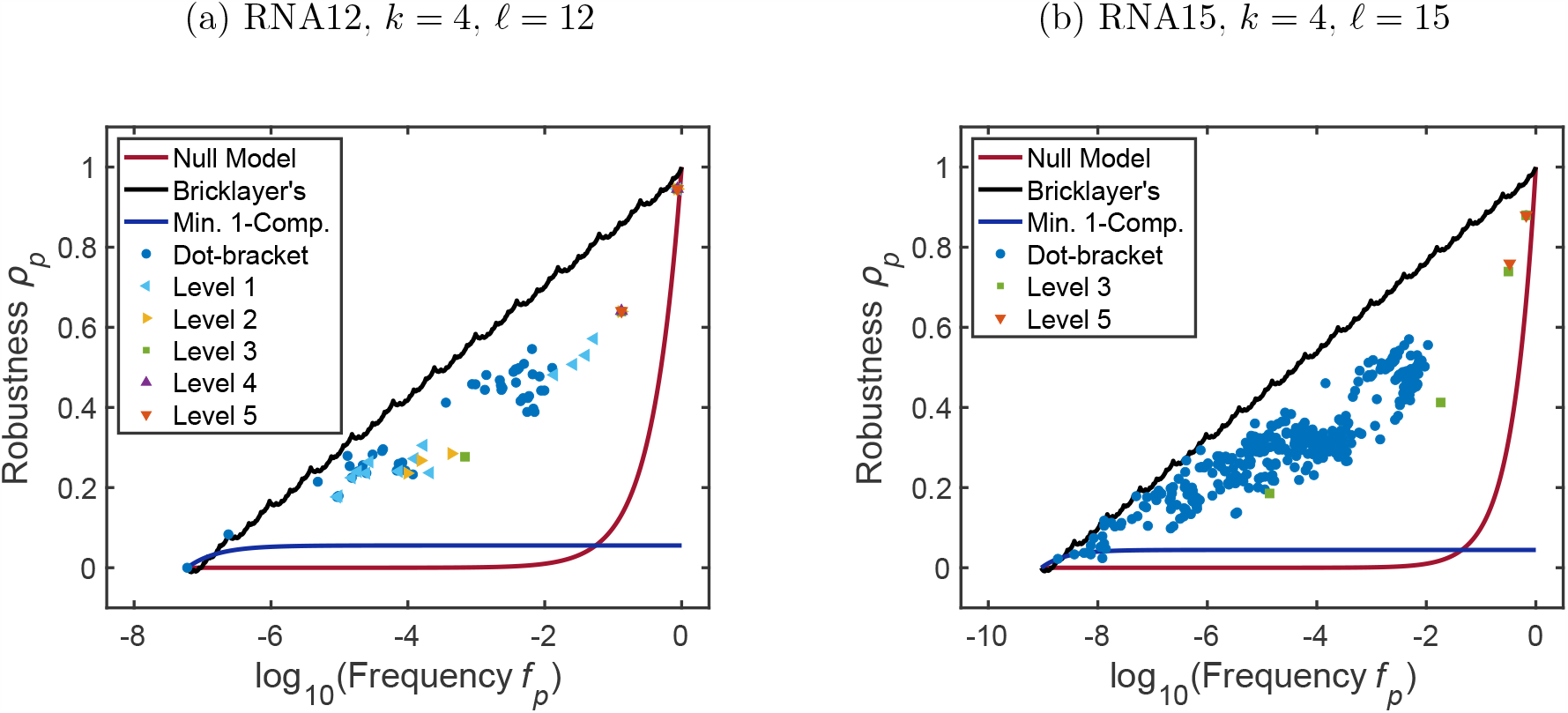
RNA abstract phenotype robustness plots for various levels of coarse-graining for (a) RNA12 and (b) RNA15 models. “Dot-bracket” structures are the standard folded RNA phenotypes which approximately match the ViennaRNA program [54] folding results. Level 1 is the first abstracted (coarse-grained) phenotype, including one or more dot-bracket structures based on coarse-grained topology. Level 2 includes phenotypes that are further coarse-grained from Level 1; Level 3 includes phenotypes that are even further coarse-grained, etc. In the case of RNA12, Levels 4 and 5 are identical because the Level 4 phenotypes are already coarse-grained as much as possible. Also plotted are the bricklayer’s bound indicating the maximum possible robustness, the null model robustness (eq. (2)), and the minimum robustness for a phenotype that contains only one component (eq. (13)); this would be the robustness of a star graph [23].

Even though the systems in Figure 6 are too small to show many Level 4 or 5 coarse-grained phenotypes, the overall trends are visible. The dot-bracket structures appear to be closest to the bricklayer’s graph maximum robustness curve. At the highest levels of coarse-graining (Level 4/5), abstract phenotypes are so densely packed with dot-bracket phenotypes that a substantial portion of the Hamming graph sequence space is covered by only a small number of abstract phenotypes. This leads to a percolation-like phenomenon that allows for highly coarse-grained, large-frequency phenotypes having high robustness as would be intuitively expected.

At lower levels of coarse-graining, however, we see a trend that we did not at first expect. It seemed to us reasonable to assume that coarse-graining dot-bracket phenotypes together, which increases the frequency, would simply “push” the robustness parallel to the diagonal logarithm line. However, the data show that coarsegrained phenotypes with sufficiently small frequencies deviate further from the maximal possible robustness (the bricklayer’s graph bound) than the phenotypes that comprise them. This is because the transition probabilities between these phenotypes being coarse-grained are likely too low to provide adequate increase in robustness after coarse-graining. Note that, in contrast to robustness, transition probabilities are expected to scale as *ϕ*_*ij*_ ≈ f_j_ for RNA [14], as well as some other models [18] such as the HP model [37] and the polyomino model for protein quaternary structure [11, 55]. In other words, the scaling of the transition probability versus frequency is no different from what would be expected from random assignment of genotype-phenotype pairs, so that these values are typically much lower than the robustness.

### B. Critical Threshold for the Coarse-Graining of Phenotypes with High Robustness

We now consider the example of coarse-graining two phenotypes and ask how much the transition probability between those phenotypes should be in order to keep them along the same diagonal robustness line parallel to the bricklayer’s graph bound. Recall that a phenotype’s neutral set *G*_*i*_ which contains *n* vertices has at most |*E*(*G*_*i*_) | = *S*_*k*_(*n*) edges, where *S*_*k*_(*n*) is once again the sums-of-digits function. We know that asymptotically *S*_*k*_(*n*) ∼ (*n*/2) log_*k*_ *n*, and a reasonable approximation to the maximum robustness is

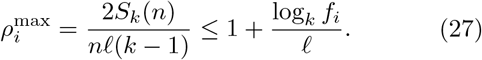

In the *high-robustness* asymptotic assumption, employed below, we assume that the robustness is near this bound and approximate it as *ρ*_*i*_ ≈ 1 + *ℓ*^−1^ log_*k*_ *f*_*i*_.

For two phenotypes *p* and *q* that are being coarse-grained into a new phenotype *S*, we can use eq. (24) to show that

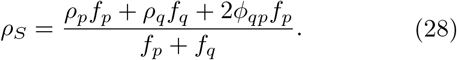

Let us assume that the two phenotypes have robustness values that are displaced from the (asymptotic) optimal robustness curve by amounts Δ_*p*_ and Δ_*q*_:

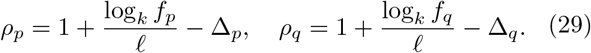

Substituting these approximations into eq. (28), we have

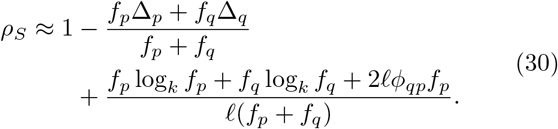

We would intuitively expect two very robust phenotypes *p* and *q* that have dense connections to each other (i.e. relatively high values of *ϕ*_*qp*_*f*_*p*_ = *ϕ*_*pq*_*f*_*q*_) to deviate from the optimal robustness curve by Δ_*p*_ and Δ_*q*_. That is to say, we expect the robustness of the coarse-grained phenotype *S* to be bounded above by

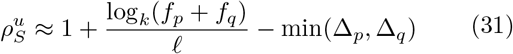

and bounded below by

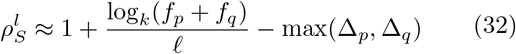

The following inequalities provide bounds on the transition probability *ϕ*_*qp*_ in order for 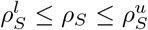 to be satisfied. These bounds are derived in Appendix B. Introducing a change of variables *β* ≡ *f*_*q*_/*f*_*p*_ and *f*_*S*_ ≡ *f*_*p*_ + *f*_*q*_ so that *f*_*p*_ = *f*_*S*_*/*(1 + *β*) and *f*_*q*_ = *f*_*S*_*β/*(1 + *β*), for Δ_*p*_ > Δ_*q*_, we have

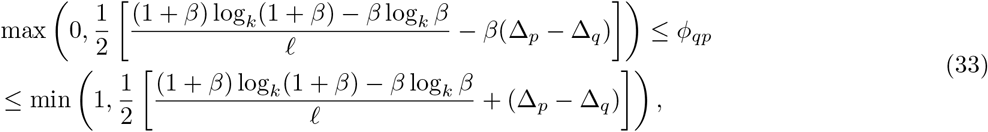

and for Δ_*p*_ < Δ_*q*_ we have

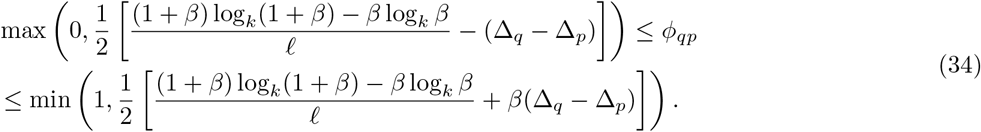

The lower and upper bounds on *ϕ*_*qp*_ for the cases Δ_*p*_ > Δ_*q*_ and Δ_*p*_ < Δ_*q*_ are plotted in Figure 7. For Δ_*p*_ = Δ_*q*_, the range 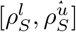 collapses to a single value, and the constraint on *ϕ*_*qp*_ becomes an equality

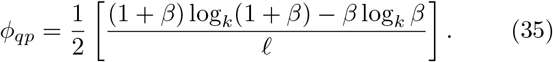

**FIG. 7.**
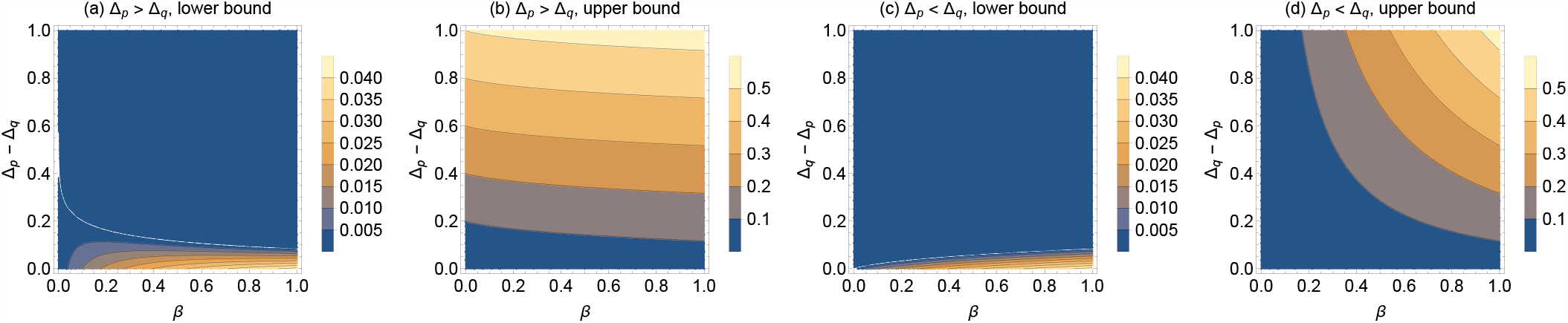
Contour plots of the (a and c) lower bounds and (b and d) upper bounds on *ϕ*_*qp*_ required for the coarse grained robustness to deviate from the maximum robustness curve by an amount that is at most max(Δ_*p*_, Δ_*q*_) and at least min(Δ_*p*_, Δ_*q*_). Contours are plotted versus the difference (a and b) Δ_*p*_ − Δ_*q*_ when Δ_*p*_ > Δ_*q*_ and (c and d) Δ_*q*_ − Δ_*p*_ when Δ_*p*_ < Δ_*q*_ on ordinate and versus the ratio of frequencies *β* ≡ *f*_*q*_/*f*_*p*_ with 0 *< β* ≤ 1 on the abscissa. None of the pairs of phenotypes from the RNA12 GP map have a transition probability which exceeds the lower bound (and therefore they are below the upper bound trivially). This suggests that maintenance of high robustness during neutral set coarse-graining requires more than two phenotypes.

We find, in fact, for any two pairs of phenotypes in the RNA12 GP map, *none* of the transition probabilities for these pairs fall within this expected range, and all of the *ϕ*_*qp*_ values are less than the theoretical lower bound required for the coarse-grained robustness to deviate from the maximum robustness no more than max(Δ_*p*_, Δ_*q*_). This makes sense when we consider the finding that for *q* ≠ *p*, in general *ϕ*_*qp*_ ≈ *f*_*q*_ [18]. While mutation of a genotype that leads to no change in phenotype is highly likely (because of high robustness), mutation to a different phenotype is equivalent to randomly picking a new phenotype, with selection probability equal to the frequency of that phenotype. The number of edges connecting two random phenotypes is too low to pass the threshold required to obtain a coarse-grained robustness that is no less than the max possible robustness minus max(Δ_*p*_, Δ_*q*_). Coarse-graining of phenotypes can, of course, bring the new robustness closer to the bricklayer’s bound in real systems, as evidenced by the most frequent Level 4 and 5 phenotypes in Figure 6. But, this is obtained because several phenotypes (not just two, the case for wich we have explicitly derived bounds) have been coarse-grained together in order to do so to maintain sufficiently high robustness. We expect that the intuition and theoretical insight gained from our formulation will be useful in the study of coarse-grained phenotypes, which is now an active area in the field of GP maps [29, 56].

## VII. DISCUSSION

In this paper we investigate maximally robust neutral sets known as bricklayer’s graphs. By applying concepts from coding theory as well as results from number theory on the sum-of-digits function, we analytically calculate the maximum phenotype robustness of a biological neutral set, a quantity which plays an important role in GP maps. We used numerical simulation to show, for the RNA sequence to secondary structure GP map and for the HP map for protein folding, that many neutral components have robustness that is near or achieves the upper bound.

We then derived a new property of sums-of-digits function and used it to calculate a lower bound on the deviation of the robustness from the maximum bound when a neutral set is made up of independent neutral components. Similar bounds for the deviation from the maximum robustness that occur when phenotypes are coarsegrained together were also derived. By coarse-graining RNA secondary structures into abstract shapes [50], we demonstrated that our bounds provide intuition for trends observed in the behaviour of the robustness of coarse-grained abstract RNA shapes.

It remains an open question as to why GP maps generically have such high robustness. There are heuristic arguments based on constrained and unconstrained sites that help point in this direction [16, 19, 57] for systems such as RNA, and it would be interesting to explore how they link to our graph-theoretical approach. The bricklayer’s graph involves genotype networks that have precisely constrained genetic sites by construction. It is therefore perhaps surprising to see that many neutral components identified in Section IV achieve the exact bricklayer’s bound, though it is possible that some of these graphs may not exactly be bricklayer’s graphs, but rather other small graphs that obtain the same bound. While combined phenotypes and neutral sets made up of multiple neutral components cannot exactly achieve this bound, it remains the case that their robustness is much closer (on a log scale) to this bricklayer’s bound than it is to a random-null model of uncorrelated phenotypes. Important ideas for future work include studying other GP maps, to see how close their robustness is to our theoretical maximum. In naturally occurring functional RNA the mutational robustness is very close to that predicted by random sampling of genotypes for the GP map [24, 33] which provides a neat example of detailed mathematical structure of the GP map being reflected in the living world. It would be extremely interesting to see if other biological systems exhibit mutational robustness that can also be predicted in this way.

Arguments from algorithmic information theory suggest that biological GP maps and other input-output maps share common underlying principles of organisation [58]. This begs the question of how close the robustnesses of non-biological systems such as spin glasses [23], quantum circuits [59], and linear genetic programs [22], are to the maximum robustness we calculate here.

Another interesting direction of future work would be to better understand the spectral properties of bricklayer’s graphs. These may provide insight into population distributions and average robustness on long evolutionary time-scales [36, 38], allowing the exploration of relationships between mutational robustness and spectral properties.

## VIII. DATA AVAILABILITY

We have introduced the web tool RoBound Calculator, a Google Colaboratory notebook which can generate, for specified *ℓ* and *k*, a continuous interpolation of the maximum robustness curve, tight upper and lower bounds on the maximum robustness curve, the exact robustnesses of bricklayer’s graphs comprised of 1 to *k*^*ℓ*^ genotypes, the random null expectation of robustness, and the minimum robustness curve for a single neutral component. The RoBound Calculator is available free of charge, with open-source code at GitHub link in ref. [42].

## IX. DECLARATION OF INTERESTS

The authors declare no competing interests.

## X. ACKNOWLEDGEMENTS

The authors thank Nora Martin and Akshay Jaggi for helpful discussions. V.M. was supported by a Marshall Scholarship and by award numbers T32GM007753 and T32GM144273 from the National Institute of General Medical Sciences. The content is solely the responsibility of the authors and does not necessarily represent the official views of the National Institute of General Medical Sciences, National Institutes of Health, or the Marshall Aid Commemoration Commission. S.N. was supported by an NSF Graduate Fellowship, a Simons Investigator award, the NSF TRIPODS program, and a Google Fellowship. S.F.G. was supported by the Engineering and Physical Sciences Research Council. S.E.A. was supported by the Royal Society and the Gatsby Foundation.

## Appendix A: Proofs of Main Text Theorems

### Theorem III.1.

*A bricklayer’s graph G*_*n,k*_(*V, E*) *with n vertices has* 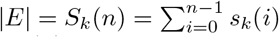 *edges, where s*_*k*_(*i*) *is the sum of all digits in the base-k representation of the integer i, and S*_*k*_(*n*) *is the sums-of-digits function*.

*Proof*. To see this, let *ℓ* be the length of the input sequence so *ℓ* ≥ log_*k*_ *n*, and let (*x*_*ℓ*−1_(*n*), …, *x*_0_(*n*)) be the vector of integers containing the digits of the base-*k* representation of the integer *n* such that 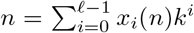. Consider the bricklayer’s graph *G*_*n*−1,*k*_. When we add one more vertex such that *G*_*n*−1,*k*_ → G_n,*k*_, we look at the base-*k* representation of *n*. An edge can be added if the base-*k* representation of *n* differs from the base-*k* representation of the neighbouring vertex by exactly one digit. Going through digit by digit, we see that the only allowed flips for the *i-*th digits are *x*_*i*_(*n*) → {0, …, *x*_*i*_(*n*) − 1}. This set has cardinality *x*_*i*_(*n*). Summing this over all digits, we find that the number of edges added to the graph *G*_*n*−1,*k*_ when adding an additional vertex *n* is the sum of digits of *n* in the base-*k* representation *s*_*k*_(*n*). Therefore, the total number of edges in *G*_*n,k*_ is 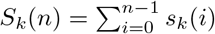. □

### Theorem V.1.

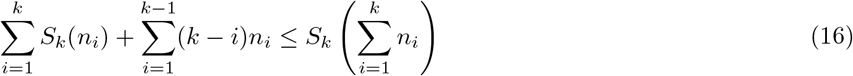

*Proof*. Let 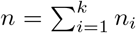, and choose *ℓ* such that *n*_*k*_ ≤ *k*^*ℓ*^. We must necessarily have that *n* ≤ *kn*_*k*_ ≤ *k*^*ℓ*+1^. Consider the Hamming graph *H*_*ℓ*+1,*k*_. Since *H*_*ℓ*+1,*k*_ = *H*_*ℓ,k*_□*K*_*k*_, we can decompose the edge set of *H*_*ℓ*+1,*k*_ into

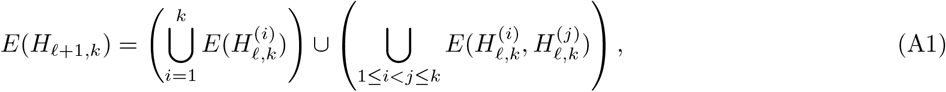

where 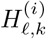 is the Hamming graph consisting of the base-*k* representations of the integers whose (arbitrarily) first digit is *i* − 1, and *E*(*G*_1_, *G*_2_) = {{*u, v*} | *u* ∈ *E*(*G*_1_) ∧ *v* ∈ *E*(*G*_2_)} is the set of edges in *H*_*ℓ*+1,*k*_ which join two subgraphs *G*_1_ and *G*_2_. Note that for 1 ≤ *i* ≤ *k*, we can construct a bricklayer’s graph 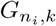 with *n*_*i*_ vertices that is a subgraph of the Hamming graph 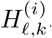. By Theorem III.1, each bricklayer’s graph has size (number of edges) equal to *S*_*k*_(*n*_*i*_). Let us assume that each bricklayer’s graph has been constructed starting on the vertex such that its index’s first digit is *i* − 1, and all other digits are 0. This ensures that the 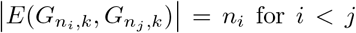 is maximal. The total number of edges in this graph *G*—i.e. the subgraph *G* of the Hamming graph *H*_*ℓ*+1,*k*_ induced by the vertex set 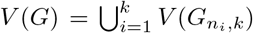—is given by the contributions from within the bricklayer’s graphs and the connections between them:

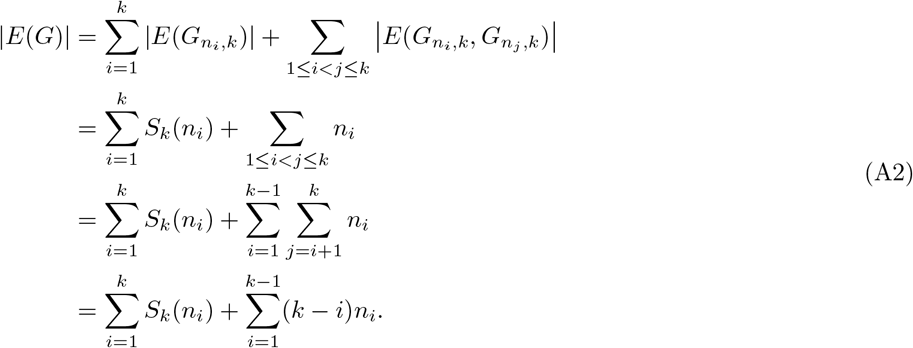

However, we also know that *G* has |*V* (*G*)| = *n* and, by Theorem III.1, size |*E*(*G*)| ≤ *S*_*k*_(*n*). Therefore,

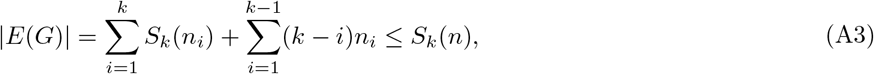

and this completes the proof. □

## Appendix B: Derivation of Bounds on Transition Probability for Coarse-Graining of Robustness

Starting with eq. (30), we now perform a change of variables to *β* ≡ *f*_*q*_/*f*_*p*_ and *f*_*S*_ ≡ *f*_*p*_ + *f*_*q*_, so *f*_*p*_ = *f*_*S*_*/*(1 + *β*) and *f*_*q*_ = *f*_*S*_*β/*(1 + *β*). Without loss of generality, we can choose phenotype assignment such that *f*_*q*_ ≤ *f*_*p*_, i.e. 0 ≤ *β* ≤ 1. We now rewrite eq. (30) as

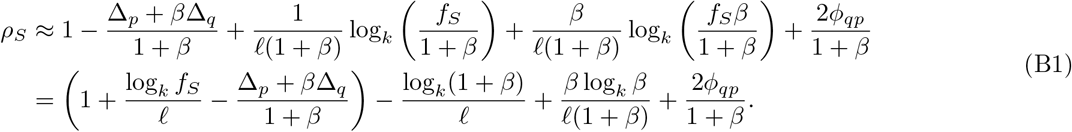

The actual coarse-grained robustness deviates from the upper bound eq. (31) by

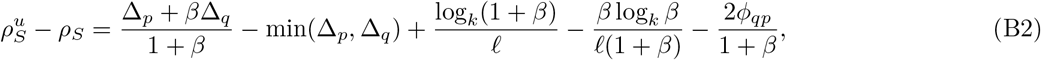

and it deviates from the lower bound by

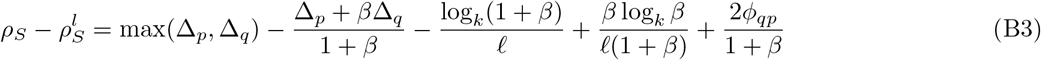

In order for the actual coarse-grained robustness *ρ*_*S*_ to fall within the bounds of the interval 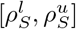, the following inequality on *ϕ*_*qp*_ needs to be satisfied, as well as the constraint 0 ≤ *ϕ*_*qp*_ ≤ 1:

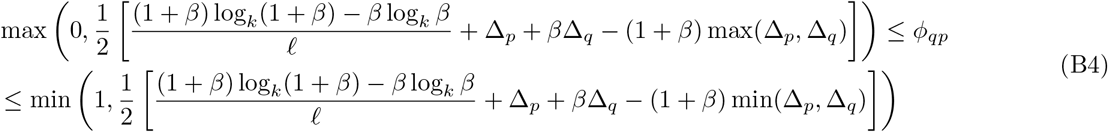

## Appendix C: Example Plots Using the RhoBound Calculator Web Tool

In this section of the Appendix, we show representative plots generated from our web tool RoBound Calculator [42], which generates—provided *ℓ, k*, and a resolution value—the continuous interpolation of the maximum robustness curve (bricklayer’s bound/blancmange-like curve), upper and lower bounds on the maximum robustness, the random null expectation robustness, the single neutral component minimum robustness, and, if desired, the exact robustnesses of bricklayer’s graphs with an integer-valued number of nodes. The examples we provide are for sequences lengths *ℓ* = 10, 30, 50, and 100 for alphabet sizes *k* = 2 (corresponding to the HP lattice protein model, in Figure 8), *k* = 4 (corresponding to the RNA alphabet, in Figure 9), and *k* = 20 (corresponding to the number of amino acids in proteins, in Figure 10). These plots help illustrate how for larger *k* and *ℓ*, the upper and lower bounds from Eq. (11) become tighter, so that the simple 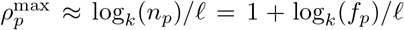 form provides good approximation to the maximum robustness.

**FIG. 8.**
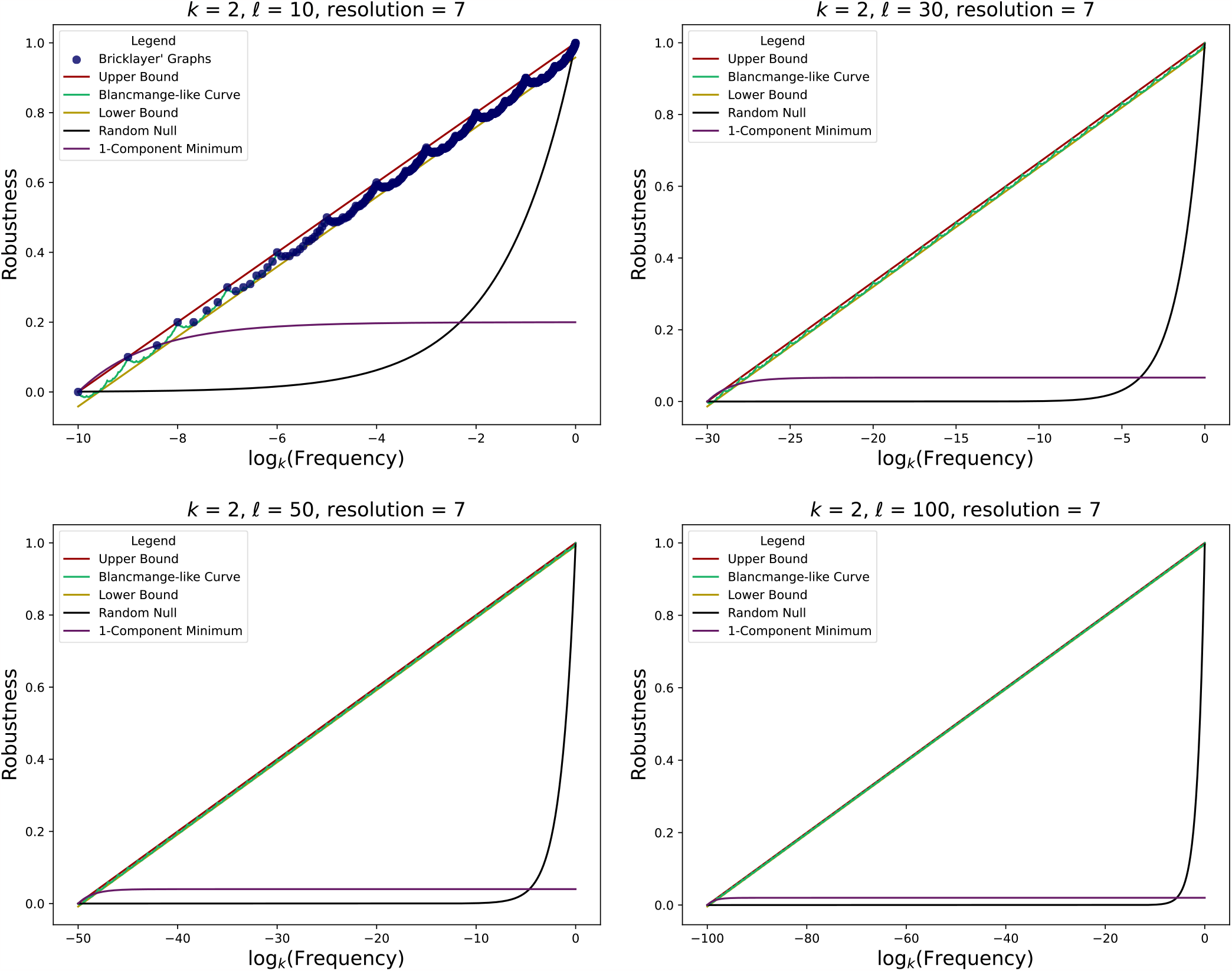
Plots of the continuous interpolation of the maximum robustness curve (bricklayer’s bound/blancmange-like curve), upper and lower bounds on the maximum robustness, the random null expectation robustness, and the single neutral component minimum robustness, generated using RoBound Calculator [42] using *k* = 2 (corresponding to a binary alphabet, such as in the HP lattice protein model). Plots for *ℓ* = 10, 30, 50, and 100 are shown. The *k* = 2, *ℓ* = 10 plot also shows, as blue dots, the exact robustness of bricklayer’s graphs with an integer number of genotypes/vertices.

**FIG. 9.**
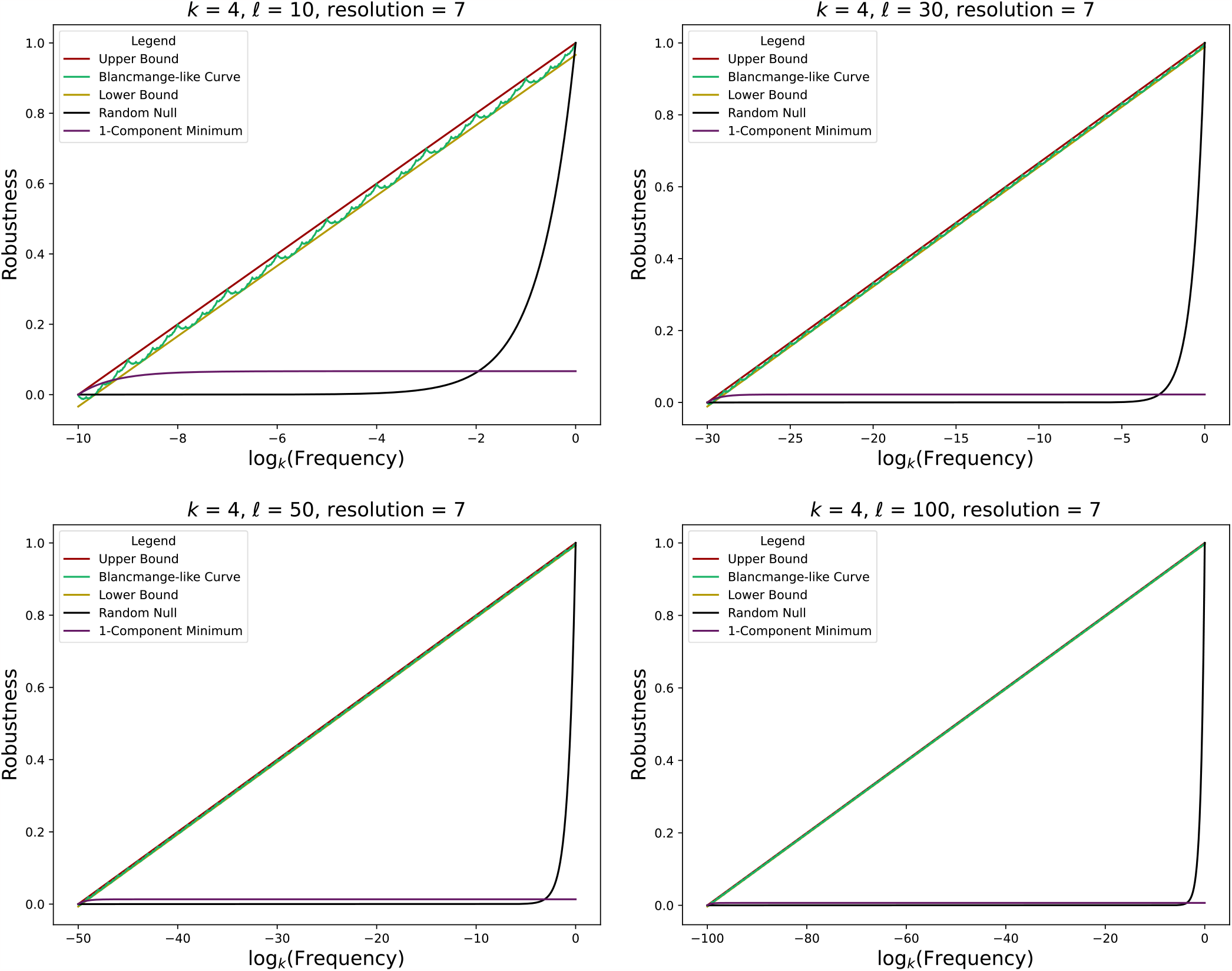
Plots of the continuous interpolation of the maximum robustness curve (bricklayer’s bound/blancmange-like curve), upper and lower bounds on the maximum robustness, the random null expectation robustness, and the single neutral component minimum robustness, generated using RoBound Calculator [42] using *k* = 4 (corresponding to the RNA alphabet). Plots for *ℓ* = 10, 30, 50, and 100 are shown.

**FIG. 10.**
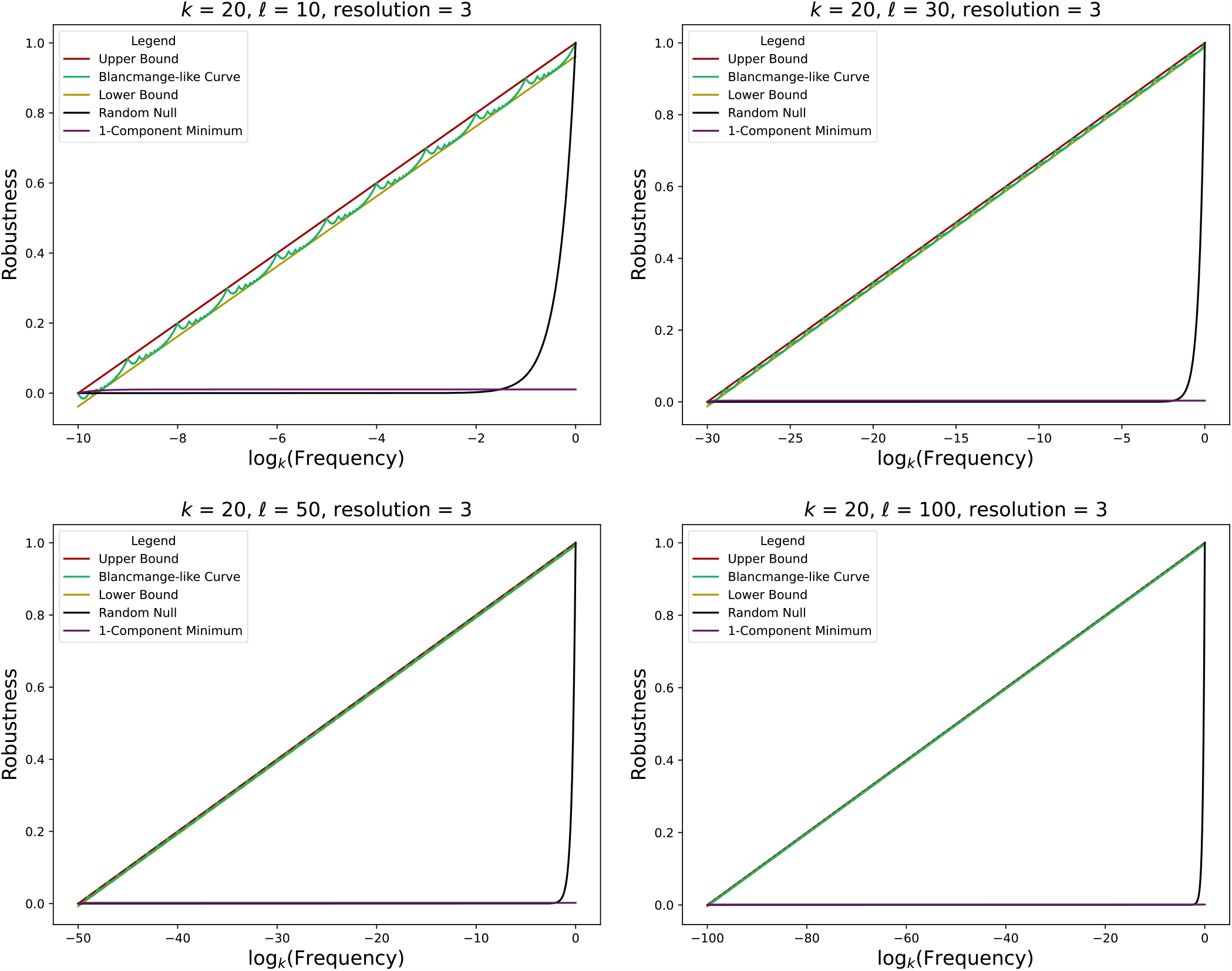
Plots of the continuous interpolation of the maximum robustness curve (bricklayer’s bound/blancmange-like curve), upper and lower bounds on the maximum robustness, the random null expectation robustness, and the single neutral component minimum robustness, generated using RoBound Calculator [42] using *k* = 20 (corresponding to the number of standard amino acids found in proteins). Plots for *ℓ* = 10, 30, 50, and 100 are shown.

## Notes

https://github.com/vaibhav-mohanty/RoBound-Calculator/

### Competing Interest Statement

The authors have declared no competing interest.

